# Deconvolution and phylogeny inference of structural variations in tumor genomic samples

**DOI:** 10.1101/257014

**Authors:** Jesse Eaton, Jingyi Wang, Russell Schwartz

## Abstract

Phylogenetic reconstruction of tumor evolution has emerged as a crucial tool for making sense of the complexity of emerging cancer genomic data sets. Despite the growing use of phylogenetics in cancer studies, though, the field has only slowly adapted to many ways that tumor evolution differs from classic species evolution. One crucial question in that regard is how to handle inference of structural variations (SVs), which are a major mechanism of evolution in cancers but have been largely neglected in tumor phylogenetics to date, in part due to the challenges of reliably detecting and typing SVs and interpreting them phylogenetically. We present a novel method for reconstructing evolutionary trajectories of SVs from bulk whole-genome sequence data via joint deconvolution and phylogenetics, to infer clonal subpopulations and reconstruct their ancestry. We establish a novel likelihood model for joint deconvolution and phylogenetic inference on bulk SV data and formulate an associated optimization algorithm. We demonstrate the approach to be efficient and accurate for realistic scenarios of SV mutation on simulated data. Application to breast cancer genomic data from The Cancer Genome Atlas (TCGA) shows it to be practical and effective at reconstructing features of SV-driven evolution in single tumors. All code can be found at https://github.com/jaebird123/tusv

## 1 Introduction

Genomic methods have provided a wealth of information about mutational landscapes of developing cancers, but have also created a great need for sophisticated computational models to make sense of the resulting data. They have revealed extensive variation patient-to-patient (intertumor heterogeneity) as well as cell-to-cell within single patients (intratumor heterogeneity) [17] and suggested a far more complex landscape of somatic variations in cancer development than earlier mutational models [20, 11] had anticipated. Extracting meaningful biological insight from such data nonetheless remains challenging. Much effort has focused on the difficulty of identifying those variants relevant to tumorigenesis and progression, known as the drivers, from the background noise of the many more chance mutations carried along with a developing tumor despite being functional irrelevant, known as the passengers [18]. More recently, attention has shifted to understanding what one can learn even from passengers regarding how a particular tumor’s mutational spectrum [1] shapes its genome across stages of progression and how that knowledge can predict its future progression and help improve prognosis. These remain substantively unsolved problems that must be better tackled if cancer researchers are to make sense of enormous and ever-growing libraries of genetic variations in cancers.

One key advance in understanding tumor genomic data was the advent of tumor phylogenetics, i.e., the use of phylogenetic inference to reconstruct tumor progression. This field arose from the observation that cancer progression is fundamentally the evolution of clonal cell populations and thus in principle interpretable via algorithms for reconstructing evolutionary trees, i.e., phylogenetics. Tumor phylogenetics itself has greatly evolved, from its initial use in making sense of intertumor heterogeneity via oncogenetic tree models [7], through the advent of methods for interpreting variation between distinct tumor regions [15, 14], between distinct cells in single tumors [22], and ultimately to recent variants that seek to explain whole-genome evolution of numerous single-cells per tumor [24, 12, 33]. Single-cell genomic data is beginning to become available in quantity, though, most studies of non-trivial patient populations are still limited to bulk sequence data, providing at best variant frequencies averaged across many single cells. Modern methods for working with such data combine phylogenetic inference with a deconvolution step, in which one infers clonal subpopulations from mixed genomic samples prior to or concurrent with inferring phylogenetic relationships between those subpopulations [27]. Numerous tumor phylogeny methods now work on this basic model of joint deconvolution and phylogenetics, with prominent examples including THeTA [21], Pyclone [25], Canopy [13], PhyloWGS [6], SPRUCE [9], and CITUP [16]. See [2, 26] for recent reviews.

Despite many advances, though, key aspects of the problem of reconstructing tumor evolution from variation data remain unresolved, an important one being the interpretation of structural variations (SVs). SVs, along with the copy number aberrations (CNAs) they frequently induce, are the primary mechanism of phenotypic adaptation in developing cancers [32]. Most tumor phylogeny methods until recently focused primarily on single nucleotide variations (SNVs) (e.g., [10, 23]). SNVs are generally abundant and make for computationally simpler analyses than other marker types but omit much of the functional mutation that we often seek to understand with tumor phylogenetics. Some early methods did focus primarily on CNAs for deconvolution [29] and phylogenetics [22, 3, 28], and several tools are now available for joint inference of SNVs and CNAs (e.g., [6, 13, 9]). There is, to our knowledge, however, no method that handles phylogenetics of SVs more comprehensively. Despite their importance, SVs introduce a number of technical challenges, including difficulty of reliable detection leading to a high expected missing data rate, of reconstructing variants that by their nature are associated with copy number variant regions of the genome, and of interpreting these more complicated event types phylogenetically.

The goal of this paper is to address the lack of methods for tumor deconvolution and phylogenetics of diverse classes of SVs at nucleotide resolution. Specifically, we develop a new method for simultaneously deconvolving inferences of SVs, derived from the Weaver variant caller [30], and reconstructing the likely evolution of clonal populations via these SV events. The method relies on a novel model extending prior literature on SNV and CNA phylogenetics [9] to handle SVs. It depends on a model of joint likelihood of genomic sequence data and clonal phylogenies, which we pose and solve through a combinatorial coordinate descent inference strategy. We demonstrate, on simulated and Cancer Genome Atlas (TCGA [19]) samples that these methods are practical and effective in inferring progression of major clones from bulk whole genome sequence (WGS) data.

## 2 Methods

### 2.1 Breakpoint and Structural Variant Definitions

Let `chrm:pos` denote the position and chromosome for each base pair in a reference genome. For example, `7:501` represents the base pair at position `501` on chromosome `7`. We define a **breakpoint** as any base pair `c:i` that is found nonadjacent to either base pair `c:i-1` or base pair `c:i+1`. If base pair `c:i` was found nonadjacent to base pair `c:i-1` we denote the breakpoint as `[c:i[` as the intact chromatin extends to the right while if base pair `c`:i was found nonadjacent to base pair `c:i+1` we denote the breakpoint as `]c:i]`. We define a **structural variant** (SV) as a pair of breakpoints found adjacent to one another in the cancer genome but at non-adjacent positions in a reference genome. We call each such pair of breakpoints a mated pair, or mates for short. For example, SV `]2:30],[5:10[` means that the segment on the reference genome on chromosome 2 at position 30 extending to the left was found next to the segment on the reference genome on chromosome 5 at position 10 in the cancer genome. This is specifically an example of a translocation SV, as the rearrangement involves different chromosomes.

To relate SVs to CNAs, we assume the reference genome is partitioned into *r* segments, with breakpoints positioned on the ends of segments excluding ends of chromosomes. (In practice, edges of segments are not always supported by breakpoints as mated breakpoints cannot always be supported with a sufficient number of reads). Each breakpoint is found in exactly one segment. Because of this, we can define both the number of times a mated breakpoint appears in a genome (denoted *c_b_* for the copy number of breakpoint *b*) and the copy number of the segments containing each breakpoint (denoted *γ_b_* for the copy number of the segment containing breakpoint *b*). A more in depth example for the appearance and copy number of breakpoints is given in Figure 1.

**Figure 1:**
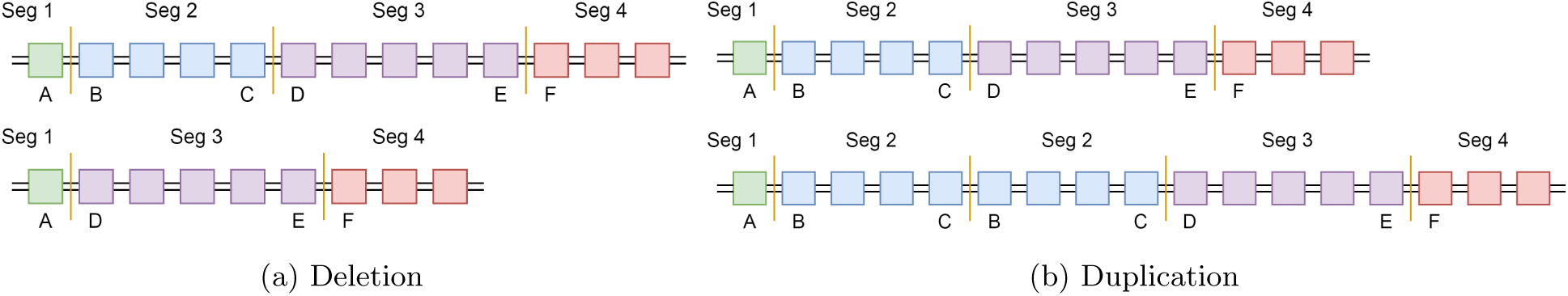
Example genomes before and after segmental deletion and duplication. Top images are the reference genome while bottom images are the genome after deletion / duplication. Each colored box represents a single base pair and base pairs between two vertical orange lines represent segments. The letters below a base pair identify the position of that base pair in the reference genome. Assume this is example holds for any single chromosome labeled `z`. (1a) shows a deletion of segment 2 (base pairs B through C) producing structural variant `]z:A],[z:D[`. The copy number of each of the mated breakpoints (*c*_]*z:A*]_ and *c*_[*z:D*[_) and the copy number of each of the segments containing these breakpoints (*γ*_]*z:A*]_ and *γ*_[*z:D*[_) are all 1 (*c*_]*z:A*]_ = *c*_[*z:D*[_ = *γ*_]*z:A*]_ = *γ*_[*z:D*[_ = 1). (1b) shows a duplication of segment 2 producing structural variant `[z:B[,]z:C]`. The copy number of each of the mated breakpoints is 1 (*c*_[z:B[_ = *c*_]z:C]_ = 1) while the copy number of each of the segments containing breakpoints is 2 (*γ*_[z:B[_ = *γ*_]z:C]_ = 2).

### 2.2 Problem Statement

Our method takes as input variant calls. We currently assume these calls are of the form produced by Weaver [30], which calls SVs and CNAs from bulk genomic read data and estimates copy numbers for copy number segments and breakpoints supporting the SVs. These variant calls are used to construct an *m* × (*ℓ* + *r*) mixed copy number matrix *F*, the rows of which represent tumor samples and columns of which represent mutations. The first *ℓ* columns correspond to breakpoints and the next *r* to mixed segmented copy numbers. The variant calls also provide a mapping of breakpoint positions to segments, which we code as an *ℓ* × *r* binary matrix *Q*. We also use information mapping breakpoints to structural variants, encoded as *ℓ* × *ℓ* binary matrix *G*. From these inputs, we seek simultaneously to infer an integer copy number matrix *C*, which describes copy numbers across the genome regions profiled for each inferred clonal cell population; a mixture fraction matrix *U*, which describes how clonal populations are distributed among tumor samples; and a phylogeny *T*, describing ancestral relationships among the clones. We assume the number of leaves *n* in the phylogenetic tree containing *N* = *2n* - 1 total nodes (clones) is known. More formally, given

> 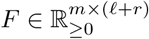 *ƒ*_*p, s*_ is the mixed copy number of variant *s* in sample *p*
>
> *Q* ∈ {0, 1}^*ℓ* × *r*^ *q*_*b, s*_ is 1 iff breakpoint *b* is in segment *s*
>
> *G* ∈ {0, 1}^*ℓ* × *ℓ*^ *g*_*s, t*_ is 1 iff breakpoints *s* and *t* are mated pairs
>
> *n* ∈ ℕ number of leaves in the phylogenetic tree
>
> *c*_*max*_ ∈ ℕ_>2_ maximum allowed subclonal copy number for breakpoints and segments
>
> *λ*_1_ ∈ ℝ_≥0_ regularization term to weight total tree cost
>
> *λ*_2_ ∈ ℝ_≥0_ regularization term to weight breakpoint consistency cost

where 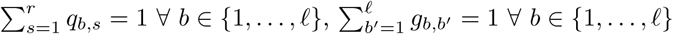, we seek to determine

> 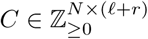 *c*_*k, s*_ is the integer copy number of segment or breakpoint *s* in clone *k*
>
> *U* ∈ [0, 1]^*m* × *N*^ *u*_*p, k*_ is the cell type *k* that makes up sample *p*
>
> *E* ∈ {0, 1}^*N* × *N*^ *e*_*i, j*_ is 1 iff directed edge (*v*_*i*_, *v*_*j*_) exists in the inferred phylogeny *T*

and minimize the objective function

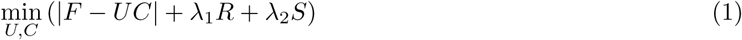

where |*F - UC*| describes the deviation between true and inferred mixed copy numbers, *R* is a phylogenetic cost, *S* is a cost capturing consistency between SVs and copy number segments, and *λ*_1_ and *λ*_2_ are regularization terms (constants).

### 2.3 Coordinate Descent Algorithm Overview

We solve for *U, C*, and *T* given *F, Q*, and *G* using coordinate descent [31]. We write two linear programs: one solving for *U* given *F* and *C* and the other solving for *C* given *U* and *F*. We then iteratively alternate between solving for *U* and for *C* while holding the other constant, either until convergence where *U* and *C* remain unchanged between iterations, or until a maximum number of iterations is reached. To avoid local minima, we run coordinate descent on multiple random initializations of *U*. Each row in *U* is independently randomly uniformly initialized so 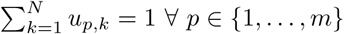 and samples independently distributed.

### 2.4 Estimating *U*

In solving for Eq. 1, we define the L1 distance |*F - UC*| as

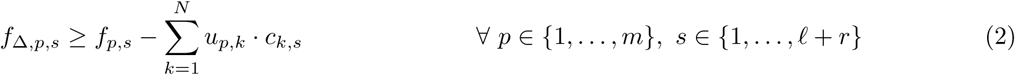

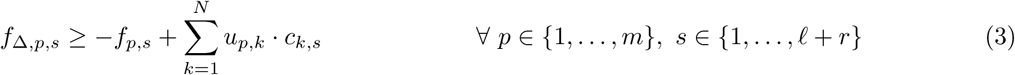

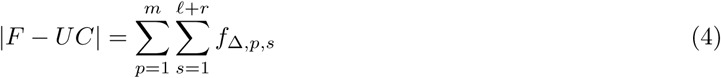

Assume then that *F* and *C* are given. To ensure each element *u*_*p, k*_ ∈ *U* is a percentage of cell type *k* in sample *p* and that percentages for a single sample sum to 1, we constrain *u*_*p, k*_ so

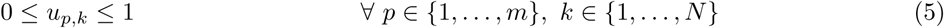

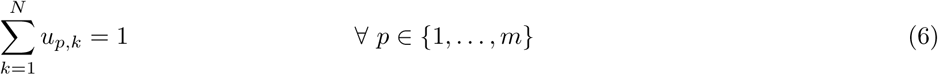

Since the regularization terms in our minimization (equation 1) do not depend on *U*, we can then simply find *U* to minimize *|F - UC|* (equation 4) given *F* and *C* subject to constraints (2), (3), (5), and (6).

### 2.5 Estimate *C*

We then estimate *C* and *T* given *F, U, Q*, and *G*.

#### 2.5.1 Binary Indicator Variables

Any variable *x* has an associated indicator variable 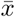 defined as

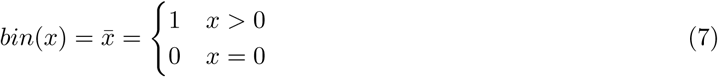

This is used throughout the following sections. To linearly define 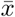, we introduce temporary variable *y*_*b*_ ∈ {0,1} as the bit representation of *x* over *q* bits [31]. The values of temporary variable *y*_*b*_ only apply to equations (8) and (9). *y*_*b*_ is then defined by

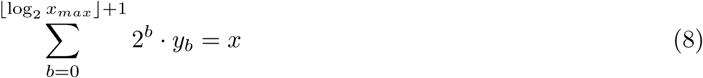

and constrains 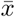 as

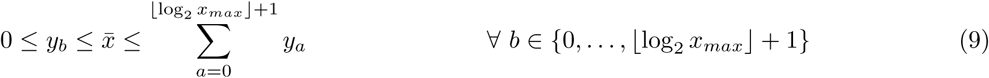

so 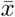 is 0 if all bits *b* are 0 and 1 if any bit of *x* is 1. In this way, any integer variable *x* with a maximum value *x*_*max*_ can be represented in binary form 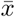. Binary indicator variables are noted with a bar on top 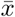 or by *bin*(*x*).

#### 2.5.2 Phylogenetic Constraints

Since the individual rows of *C* are not independent but instead share a phylogenetic history, we create a tree structure *T* representing the inferred relationships between rows in *C*. We define a binary tree *T* using a *N* × *N* directed adjacency matrix *E*. To impose a tree structure on *E*, assume the first *n* clones are leaf nodes and clones *n* + 1 through 2*n* - 1 = *N* are internal nodes, with node *N* as the root. We constrain element *e*_*i, j*_ as follows:

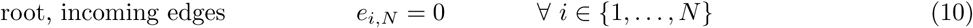

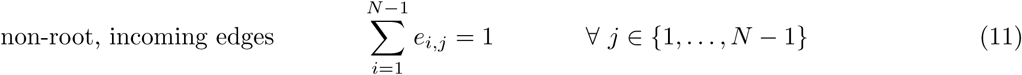

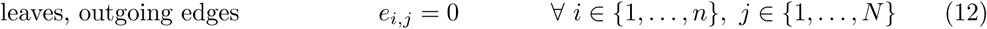

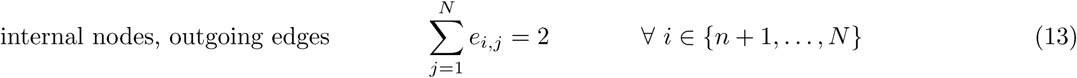

Equations (10) and (11) ensure the root has no in-edges and all other nodes have exactly one in-edge. Equations (12) and (13) force leaves to have no out-edges and all internal nodes to have exactly two out-edges.

#### 2.5.3 Phylogenetic Cost

We next ensure all copy numbers are below some input maximum *c*_*max*_ and force the normal (non tumor) root node to be diploid (each segment having copy number 2) and free of structural variants (copy number of all breakpoints is 0):

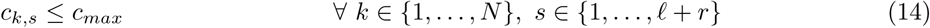

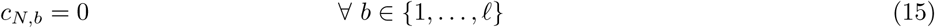

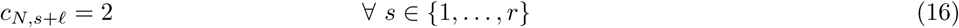

We next model a phylogenetic tree cost, using CNAs to estimate evolutionary distance *ρ*_*i, j*_ across each tree edge (*v*_*i*_, *v*_*j*_) ∈ *E*. We approximate evolutionary distance by the L1 distance between the copy number profiles of an edge’s endpoints 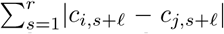. While there are more sophisticated models of copy number distance in the literature [5, 28, 4, 8], we use L1 distance as an approximation as it can be coded and computed efficiently within the ILP framework. To linearly define *ρ*_*i, j*_ we use temporary variable *x*_*i, j, s*_ ∈ ℕ^*N*×*N*×*r*^, defined as the absolute change in copy number of segment *s* on edge (*v*_*i*_, *v*_*j*_). Here, the values of temporary variable *x*_*i, j, s*_ only apply to equations (17) through (20).

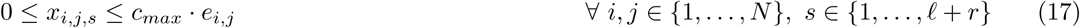

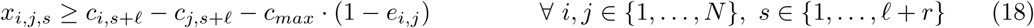

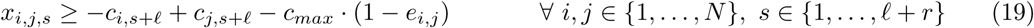

Equation (17) sets the cost to zero for any pair of nodes (*v*_*i*_, *v*_*j*_) where *v*_*i*_ is not the parent of *v*_*j*_, while equations (18) and (19) set the cost to be the absolute difference between copy number for of end nodes for any edge (*v*_*i*_, *v*_*j*_). We then define the cost across edge (*v*_*i*_, *v*_*j*_) and total cost of tree as

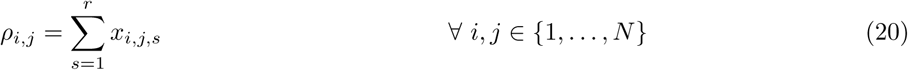

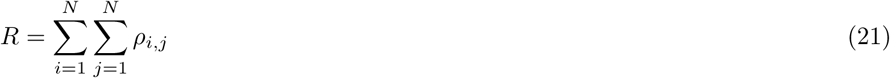

#### 2.5.4 Perfect Phylogeny on Appearance of Breakpoints

We next impose a perfect phylogeny on breakpoints. While the perfect phylogeny assumption is problematic for other variant types, we argue that it is sufficiently unlikely for a base-resolution breakpoint to recur that it can be neglected. Note that violations of the infinite site model due to allelic loss are handled separately by treating a lost allele as having copy number zero. We therefore impose constraints to force each breakpoint to appear across exactly one edge in *T* and for mated breakpoints to appear together. Define *W* ∈ {0, 1}^*N* × *N* × *ℓ*^, where each element *w*_*i, j, b*_ is 1 if the copy number of breakpoint *b* goes from 0 to a positive integer across edge (*v*_*i*_, *v*_*j*_) and 0 otherwise. To linearly define *w*_*i, j, b*_ we define temporary variable *x*_*i, j, b*_ ∈ {0,1, 2, 3} to be

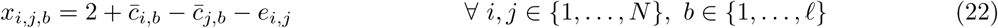

so *x*_*i, j, b*_ is 0 iff the copy number of breakpoint *b* increases from 0 across edge (*v*_*i*_, *v*_*j*_). The value of temporary variable *x*_*i, j, b*_ only applies to equations (22) and (23). Using the binary representation 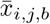 of *x*_*i, j, b*_, define *w*_*i, j, b*_ and ensure *w*_*i, j, b*_ is 1 for a single edge in the tree.

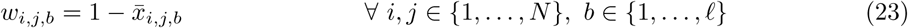

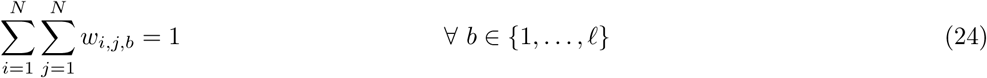

Using breakpoint mate indicator *g*_*s, t*_ ∈ {0,1}, where *g*_*s, t*_ is 1 iff breakpoints *s* and *t* are mates, we force breakpoint indicators to be equal for mates.

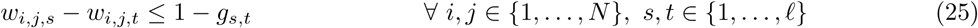

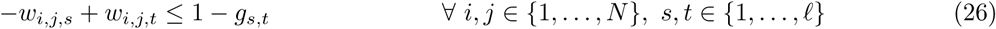

Note we extend the notation of breakpoint appearance indicator *w*_*i, j, b*_ to have 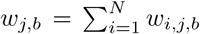 be 1 if breakpoint *b* appears at node *v*_*j*_ and 0 otherwise.

#### 2.5.5 Ancestry Condition for Non-Disappearing SVs

We next impose the two-state perfect phylogeny ancestry condition as described in [10] for the appearance of breakpoints. For any breakpoint *s* that appears as an ancestor to breakpoint *t*, the total fraction of cells with breakpoint *s* must be larger than the fraction with breakpoint *t* so long as breakpoint *s* never subsequently disappears. To enforce this, the fraction of cells *φ*_*p, b*_ containing breakpoint *b* in sample *p* is defined as

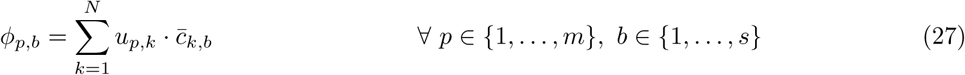

We then must define a few variables to force *φ*_*p, s*_ ≥ *φ*_*p, t*_ if breakpoint *s* appears before breakpoint *t* and is never subsequently lost. Let *v*_*i*_ be the *i*^*th*^ node in the phylogeny and *v*_*i*_ ≺ *v*_*j*_ denote that node *v*_*i*_ is an ancestor of *v*_*j*_. We first define ancestor variables *a*_*i, j*_ ∈ {0,1} as 1 if *v*_*i*_ ≺ *v*_*j*_ and 0 otherwise for all *i, j* ∈ {1, …, *N*}. Linearly define *a*_*i, j*_ by

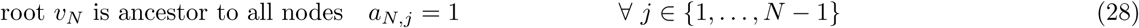

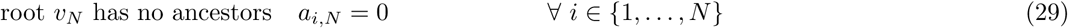

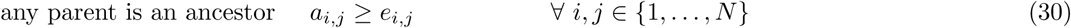

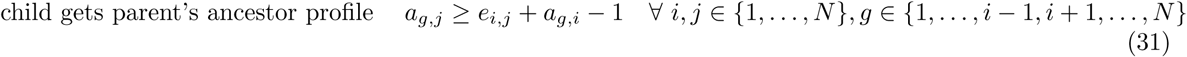

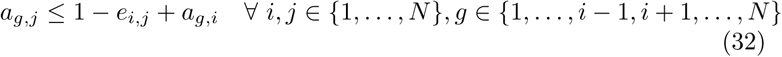

Next, define the number of descendants to node *v*_*i*_ with at least one copy of breakpoint *b* as *d*_*i, b*_ for all *i* ∈ {1, …, *N*}, *b* ∈ {1, …, *ℓ*}. To linearly define *d*_*i, b*_, define temporary binary variables *x*_*i, j, b*_ ∈ {0,1} for equations (33) through (36) for all *i, j* ∈ {1, …, *N*}, *b* ∈ {1, …, *ℓ*} to be 1 if *a*_*i, j*_ and 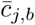 and zero otherwise.

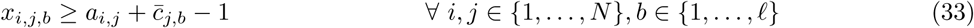

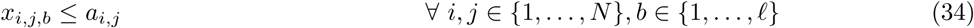

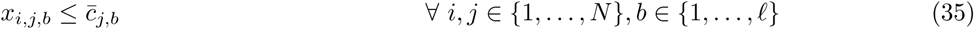

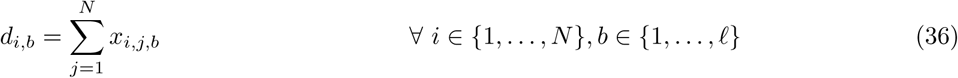

Define temporary binary variables *ȳ*_*i, b*_ ∈ {0,1} for equations (37) through (39) to be 0 to be zero if all descendants of node *v*_*i*_ contain at least one copy of breakpoint *b* and 1 otherwise.

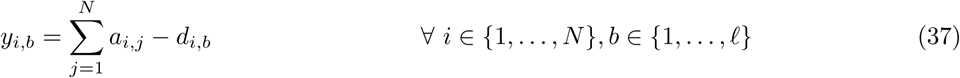

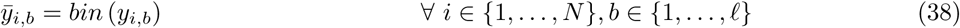

Define temporary binary variable 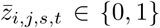 ∈ {0,1} for equations (39) and (40) to be 0 only if breakpoint *s* appears at node *v*_*i*_, breakpoint *t* appears at node *v*_*j*_, node *v*_*i*_ is an ancestor to node *v*_*j*_, and breakpoint *s* never disappears.

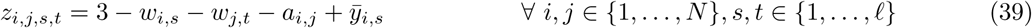

Finally, apply the condition that the fraction of cells *φ*_*p, s*_ containing breakpoint *s* in sample *p* must be larger than the fraction of cells *φ*_*p, t*_ containing breakpoint *t* in sample *p* if breakpoint *s* appears in an ancestor to the node where breakpoint *t* appears and breakpoint *s* is never lost in any descendant (no descendant has copy number 0 for breakpoint *s*).

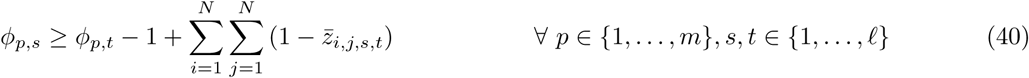

Note that 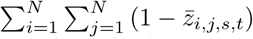 can only take on values 0 or 1 since breakpoint appearance indicator *w*_*i, s*_ and *w*_*j, s*_ can only be both 1 at most once across all *i, j*. This means the condition *φ*_*p, s*_ ≥ *φ*_*p, t*_ only holds when breakpoint *s* appears before breakpoint *t* and never subsequently disappears. Note the ancestry condition is implied by but weaker than the sum condition described in [10], but can similarly be enforced by linear constraints.

#### 2.5.6 Structural Variant and Segment Consistency

Since each breakpoint belongs to exactly one segment, we define the copy number of each segment containing a breakpoint *b* and constrain it so a breakpoint’s copy number never exceeds that of its containing segment:

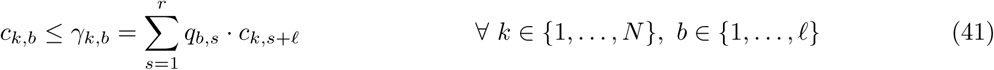

where input *q*_*b, s*_ ∈ {0,1} is 1 if segment *s* contains breakpoint *b*. 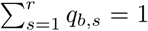 as each breakpoint belongs to a single segment. We similarly define *ψ*_*p, b*_ directly from the input to be the mixed copy number of the segment containing breakpoint *b*.

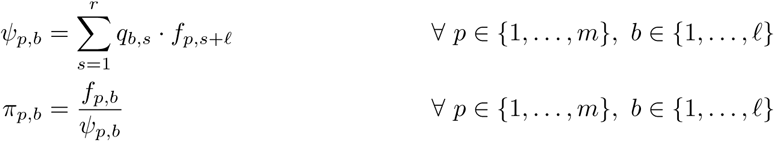

Intuitively, the ratio *π*_*p, b*_ of the mixed copy number of a breakpoint to the mixed copy number of the segment containing that breakpoint should be maintained in the integer output as this preserves the difference in mutation types (duplication, deletion). To penalize for discrepencies between the inferred ratio of breakpoint and its segment copy number given *π*_*p, b*_, we incorporate the following quantity into our objective function:

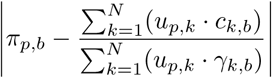

To convert this from a ratio to units of copy numbers, we rearrange the expression and define *S* for the final term in the objective function (1) to be

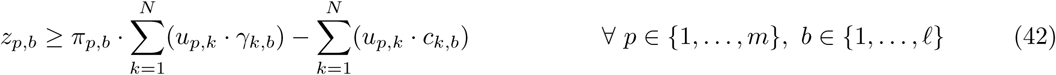

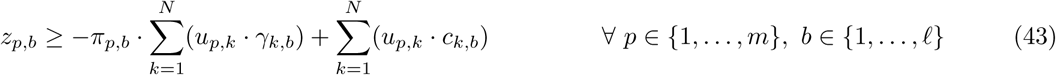

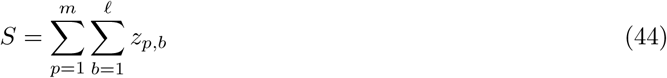

In this way, increased emphasis is placed on the relationship between segments and breakpoints. The solution for *C* and *T* is found by minimizing (1) subject to constraints (2) through (44).

## 3 Results

### 3.1 Simulated Data

To validate accuracy of the method on data of known ground truth, we assess accuracy in inference of copy number profiles across clones. For each such test, we generate a copy number matrix *C*_*tru*_ containing breakpoints and segments, mix this matrix with a mixture fraction matrix *U*_*tru*_ to get the mixed copy number matrix (*C*_*tru*_ × *U*_*tru*_ → *F*), run our deconvolution algorithm, and compare the inferred copy number matrix *C*_*in ƒ*_ with the original true copy number matrix *C*_*tru*_. We score our result as the L1 distance (|*C*_*tru*_ - *C*_*in ƒ*_|) between copy number matrices after a maximum matching between copy number profiles (for clones).

To generate *C*_*tru*_, we simulated mutation data varying the expected number of mutations l, number of samples *m*, and number cell types *n*. For each triplet (*l, m, n*), five synthetic patients were generated. Reported scores are averaged across those 5 patients. For each run of the simulation, we generated a binary tree *T* with *n* leaves and a random topology. Mutations were assigned so that the expected numbers of mutations across all edges in each tree are equal. We start with a genomic profile for the root (assumed to be a normal diploid cell containing no structural variants) and progressively added a Poisson-distributed number of mutations across each edge down to the leaves. Initially, the root node contains three pairs of homologous chromosomes of the same lengths as human chromosomes 1-3. To generate mutations, a central location is uniformly chosen across all chromosomes, then a mutation size is sampled from an exponential distribution, with expectation equal to the mean structural variant size found across 59 TCGA samples. The mutation type is uniformly randomly selected to be either a tandem duplication, deletion, or inversion. From the generated tree, we obtain a copy number matrix *C*_*tru*_. We then create a cell type mixture matrix *U*_*tru*_ by uniformly randomly assigning cell type fractions such that the fraction of all cell types in each sample sums to 1. *U*_*tru*_ and *C*_*tru*_ are subsequently multiplied to generate mixed copy number matrix *F*.

Since there is no method for validating how accurate the choice of regularization terms *λ*_1_ and *λ*_2_ are on real data, we define empirical values for these terms based on each sample and show they perform well on simulated data. We choose regularization terms *λ*_1_ and *λ*_2_ empirically from the data to be 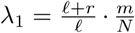 and 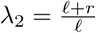. This allows the maximum error in the |*F - UC*| term in the minimization, which is *m*·(*ℓ*+*r*)·*c*_*max*_, to equal the maximum errors in *λ*_1_*R* and *λ*_2_*S* terms, which are *ℓ*·*N*·*c*_*max*_ and *ℓ*·*m*·*c*_*max*_ respectively. To show these empirical definitions do as well as iteratively choosing the hyperparameters, we test on simulated data generated for *n* = 3 leaves, *m* = 3 samples, and *l* = 50 mutations as this produces approximately 100 breakpoints, a value comparable to the average number of breakpoints found in real, TCGA samples. Figure 2 shows that emperically selecting hyperparameters *λ*_1_ and *λ*_2_ (solid green curve) performs reasonably well as compared to other hyper parameter values (dotted blue curve). Both outperform the algorithm when excluding the regularization terms (dashed red curve) indicating the usefulness of including phylogenetic cost and breakpoint-segment consistency into the model.

**Figure 2:**
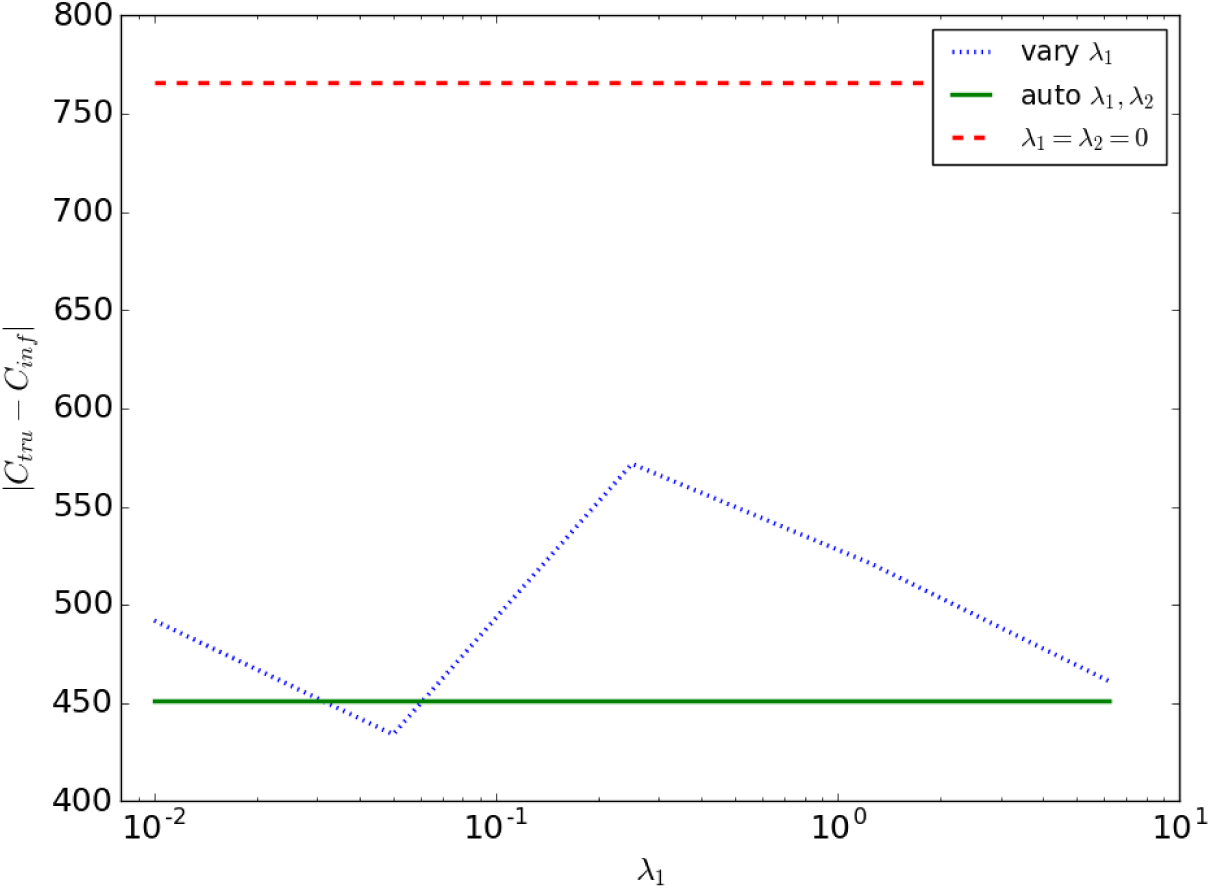
Score on simulated data for varying hyperparameters *λ*_1_ and *λ*_2_. Score is the L1 distance between the true *C_tru_* and inferred *C_in ƒ_* copy number matrices after maximum matching (lower means better performance). Dotted blue line is score when *λ*_2_ = 0 is held constant and *λ*_1_ varies from 0.01, 0.05, 0.25, 1.25, 6.25 across the x-axis. Solid green line shows the score when hyperparameters are automatically chosen based on the number of SVs and CNAs in the data set while dashed red line shows score for when only a single term in the optimization expression is used.

### 3.2 TCGA Data

We next apply the methods to a selection of TCGA breast cancer (BRCA) samples [19], restricting analysis to a subset of 59 samples for which WGS data was available. Of these, 31 ran successfully within our prescribed run time limit, while 28 with the highest SV counts timed out before completion or required more memory than was available to us (we have access to 128Gb of RAM). Since there is no known ground truth for these samples, we cannot assess their individual accuracy. Nonetheless, they provide some basis for analysis of trends across samples. Space does not permit us to display all observed trees, so for purposes of illustration we classify them into seven observed topologies (A-G), shown in Fig. 3, with frequencies of occurrence shown in Fig. 4. None of the inferred trees are purely linear, consistent with a model of significant subclonal heterogeneity rather than a simple sequential model of clonal progression. Quantitation by several measures of heterogeneity, as shown in Fig. 5, likewise suggests a wide diversity among samples. The data is suggestive of a possible clustering into distinct low-diversity and high-diversity subclusters, but with substantial overlap between clusters.

**Figure 3:**
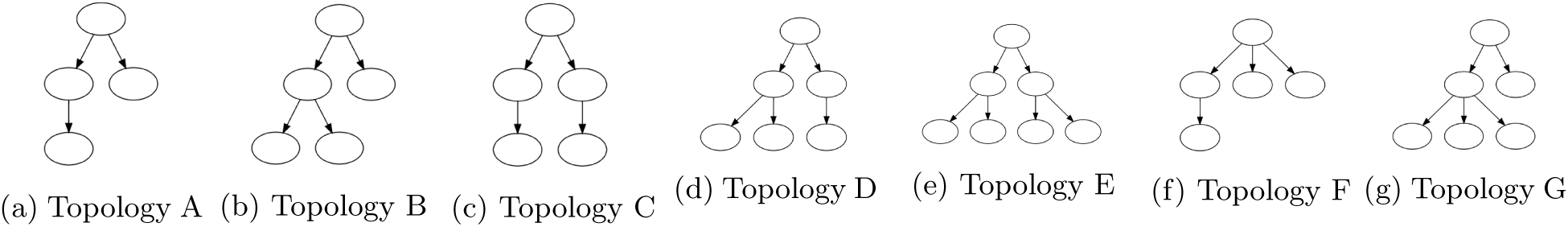
Tree topologies observed across 31 TCGA BRCA samples, grouped into seven categories (A-G).

**Figure 4:**
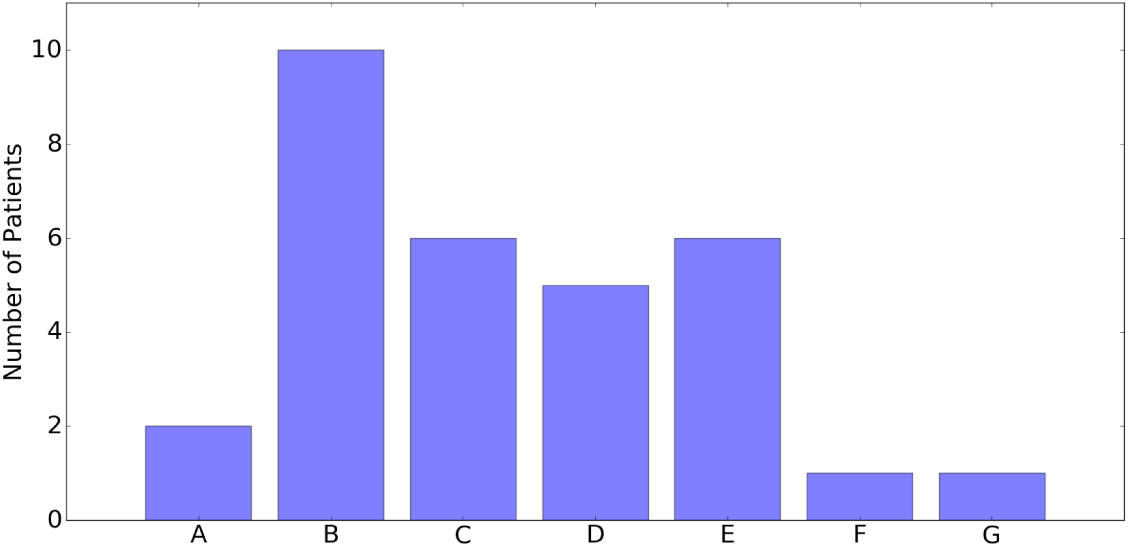
Histogram of occurrences of tree topologies across 31 TCGA BRCA samples.

**Figure 5:**
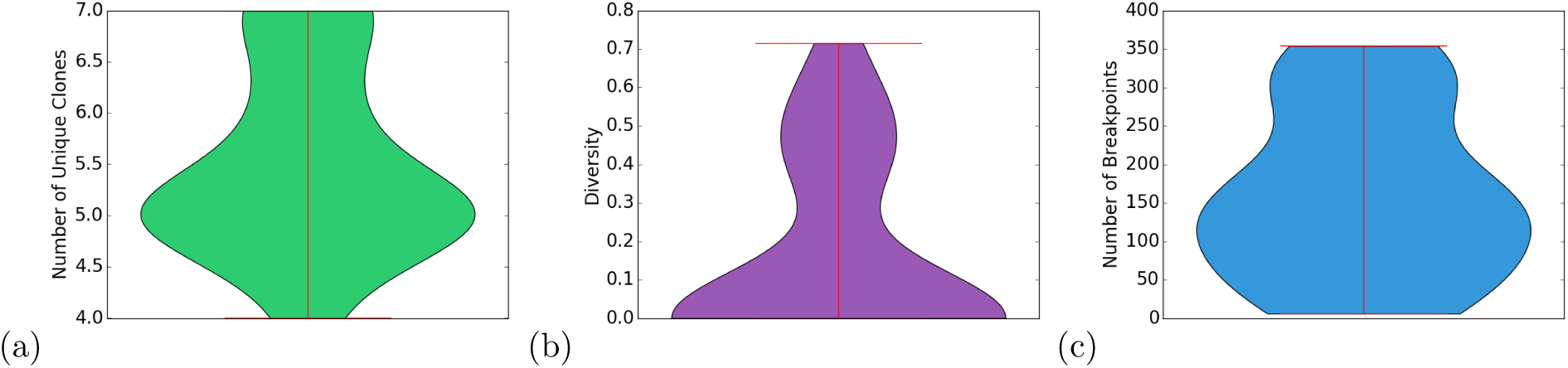
Violin plots quantifying heterogeneity of the 31 TCGA trees. (a) Number of unique clones. (b) Diversity, defined in terms of the clonal frequency vector *u*_*p, k*_ as 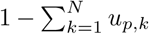 Number of breakpoints.

## 4 Discussion

We have developed a new method for automated joint deconvolution and phylogeny inference of tumor genomic data designed to address the important unsolved problem of describing progression via SVs. We specifically learn a model encompassing CNAs and SVs of major clones, mixture fractions of these clones across samples, and phylogenetic tree relating the clones. We pose the model inference problem to balance the likelihood of sequence read data with respect to copy numbers and observed breakpoints against the evolutionary cost of the phylogenetic tree. We solve the resulting model via a coordinate descent algorithm posed as a pair of MILPs. We demonstrate that the method can accurately and efficiently reconstruct clonal populations and phylogenetic histories from both simulated tumor data. Application to WGS data from the TCGA shows the method to be effective on real data supportive of a range of tree topologies and complexities.

This work provides a proof-of-concept demonstration of the feasibility of more comprehensively modeling the important role of SVs in tumor evolution, but also suggests a number of avenues for future work. Our methods currently rely on a sometimes costly and potentially suboptimal model fitting algorithm, and further algorithmic advances might plausibly lead both to greater efficiency and improved solution quality.

In particular, there are currently practical limits on the total SNV counts the method can handle without excessive run time and memory usage. While most of the TCGA breast cancers considered fell within those limits, a significant minority did not. Furthermore, our work focuses only on the subproblem of handling SVs (and associated CNAs), and will likely benefit from incorporating other variant types, most notably SNVs but also potentially expression, methylation, or other markers of cell state. Furthermore, biotechnology for data generation is continuing to advance, with growing numbers of computational methods taking advantage of single cell sequence or long read technologies that can provide direct single-cell readouts of SNV or CNA data. SV detection is problematic for all current single-cell technologies, and we can anticipate value in combining single-cell methods with bulk deconvolution methods such as ours for SVs. Finally, the present work has focused only on the development of the new technology and its validation. The ultimate value of the work will lie in bringing SV-aware phylogenetics to diverse patient cohorts, to begin to develop a comprehensive understanding of the landscape of SV variation in tumor progression and its implications for patient prognosis and treatment.

## Acknowledgments

We thank Ashok Rajaraman and Jian Ma for helpful discussions and assistance with Weaver. Portions of this work have been funded by the U.S. National Institutes of Health via award R21CA216452 and Pennsylvania Dept. of Health Grant GBMF4554 4100070287. The Pennsylvania Department of Health specifically disclaims responsibility for any analyses, interpretations or conclusions. This work used the Extreme Science and Engineering Discovery Environment (XSEDE), which is supported by National Science Foundation grant number OCI-1053575. Specifically, it used the Bridges system, which is supported by NSF award number ACI-1445606, at the Pittsburgh Supercomputing Center (PSC). The results published here are in whole or part based upon data generated by The Cancer Genome Atlas managed by the NCI and NHGRI. Information about TCGA can be found at http://cancergenome.nih.gov.

